# Single-Nucleus Chromatin Accessibility Identifies a Critical Role for TWIST1 in Idiopathic Pulmonary Fibrosis Myofibroblast Activity

**DOI:** 10.1101/2022.01.10.475117

**Authors:** Eleanor Valenzi, Harinath Bahudhanapati, Jiangning Tan, Tracy Tabib, Daniel I. Sullivan, John Sembrat, Li Fan, Kong Chen, Mauricio Rojas, Audrey Lafargue, Dean W. Felsher, Phuoc T. Tran, Daniel J. Kass, Robert Lafyatis

**Author notes:** denotes equal contribution (Daniel J. Kass, Robert Lafyatis). Corresponding Author: Eleanor Valenzi, MD, University of Pittsburgh, NW628, UPMC Montefiore, 3459 Fifth Avenue, Pittsburgh, PA 15213.

## Abstract

In idiopathic pulmonary fibrosis (IPF) myofibroblasts are key effectors of fibrosis and architectural distortion by excessive deposition of extracellular matrix and their acquired contractile capacity. Single-cell RNA-sequencing has precisely defined the IPF myofibroblast transcriptome, but identifying critical transcription factor activity by this approach is imprecise. We performed and integrated snATAC-seq and scRNA-seq from human IPF and donor control explants to identify differentially accessible chromatin regions and enriched transcription factor motifs within lung cell populations. TWIST1 and other E-box transcription factor motifs were significantly enriched in IPF myofibroblasts compared to both IPF non-myogenic and control fibroblasts. *TWIST1* expression was also selectively upregulated in IPF myofibroblasts. Overexpression of Twist1 in lung COL1A2-expressing fibroblasts in bleomycin-injured mice was associated with increased collagen synthesis. Our studies utilizing human multiomic single-cell analyses combined with *in vivo* murine disease models confirm a critical regulatory function for TWIST1 in IPF myofibroblast activity in the fibrotic lung. Understanding the global process of opening TWIST1 and other E-box TF motifs that govern myofibroblast differentiation may identify new therapeutic interventions for fibrotic pulmonary diseases.

## Introduction

Idiopathic pulmonary fibrosis (IPF) is a devastating, aging-associated fibrotic lung disease resulting in tissue architectural distortion and impaired gas exchange, ultimately progressing to respiratory failure and death in most patients. Current therapeutics have limited effect, and there are no approved medications that convincingly improve mortality or quality of life (1). While the precise pathogenesis remains unknown, current paradigms propose that repetitive microinjuries of the alveolar epithelium provoke dysregulated crosstalk with the mesenchymal compartment leading to expansion of an activated myofibroblast population within so-called fibroblastic foci (2). Myofibroblasts are key effector cells of fibrosis by excessive deposition of extracellular matrix proteins and by their acquired contractile capacity resulting in distorted lung architecture (3). In IPF myofibroblasts are also resistant to apoptosis, overcoming their normal clearance mechanisms that occur during physiological regeneration such as wound healing (4). Myofibroblasts are the primary collagen-producing cell propagating fibrosis in diverse organs, with a high disease burden ranging from the dermal and lung fibrosis of systemic sclerosis, to nephrogenic fibrosis, cirrhosis, and graft versus host disease (5).

In recent years, the widespread adoption of single-cell RNA-sequencing (scRNA-seq) has produced multiple large cell atlases of the human control and IPF lung, allowing for precise characterization of the myofibroblast and other cell population transcriptomes (6–9). While gene expression provides critical information on defining a cell’s phenotype and active signaling pathways, defining upstream gene regulatory networks from the transcriptome alone is imprecise. Temporal control of gene expression is regulated by the cooperative interactions of *trans-*acting DNA binding proteins with *cis-*regulatory element regions within the genome, such as promoters and enhancers (10). These sequences dictate expression of target genes in a cell-type dependent manner by recruiting sequence-specific transcription factors (TFs) (11). The advent of single nucleus assay for transposase-accessible chromatin sequencing (snATAC-seq) technology now allows the study of open chromatin regions in heterogenous cell populations, thus connecting the input regulatory signals with the output gene expression defining each cell population and its effector phenotype (12). Given the central role for myofibroblasts in the pathogenesis of pulmonary fibrosis and our current lack of any drug targeting these cells, delineating their TF regulation and chromatin accessibility may identify new targets for IPF and other fibroproliferative disorders.

Here we integrated snATAC-seq and scRNA-seq from human IPF and donor control explants to identify differentially accessible chromatin regions and transcription factor motifs (consensus-sequence specific binding sites) within human lung cell populations. We specifically focus on IPF myofibroblasts and identify enrichment of E-box TF motifs in IPF myofibroblasts compared to both IPF non-myogenic and control fibroblasts. As the TF TWIST1 is selectively expressed in IPF myofibroblasts, it was particularly implicated as positive putative regulator of IPF myofibroblast differentiation. We investigated Twist1 *in vitro* and in an animal model of pulmonary fibrosis. Our studies demonstrate a critical role for TWIST1 in regulating myofibroblast effector functions in IPF.

## Results

### Single-cell transcriptional and chromatin accessibility profiling in the IPF and donor control lung

We performed scRNA-seq and snATAC-seq on three IPF and two donor control lung tissue samples (Figure 1A, B) with 8,738 nuclei included for snATAC-seq analyses after filtering. Subpleural lower lobe tissue was collected at the time of lung transplant in individuals with IPF, and from organ donors with no preexisting lung disease whose lungs were unable to be matched for transplant. Histologic review of adjacent tissue showed usual interstitial pneumonia for all IPF samples (Supp Fig 1). Each sample was processed immediately upon collection from the operating room. To increase the robustness of our scRNA-seq dataset, we included 13 other scRNA-seq samples processed by our laboratory for a total of 18 samples (n=10 IPF, 8 Control) with 65,179 cells included after filtering (Figure 1C, D). Batch correction was performed by reciprocal principal component analysis (PCA) integration for the scRNA-seq dataset, and with the R package Harmony for the snATAC-seq dataset (13). The R packages Seurat and Signac were used for dimensional reduction, clustering, differential expression/accessibility testing, and visualization at the individual sample and aggregate dataset level (14, 15). We annotated snATAC-seq cell types by transferring predicted labels (Supp Fig 3A) of the scRNA-seq dataset based on their transcriptional profiles (Figure 1D), in addition to manually identifying nuclei cell types by examining chromatin accessibility matrices for cell type marker gene activity (a measure of chromatin accessibility within the promoter and gene). Comparison between snATAC-seq cell-type predictions by label transfer and manual annotations of the unsupervised clustering indicated that all major cell types were present in both datasets and consistently identified by both methods.

**Figure 1.**
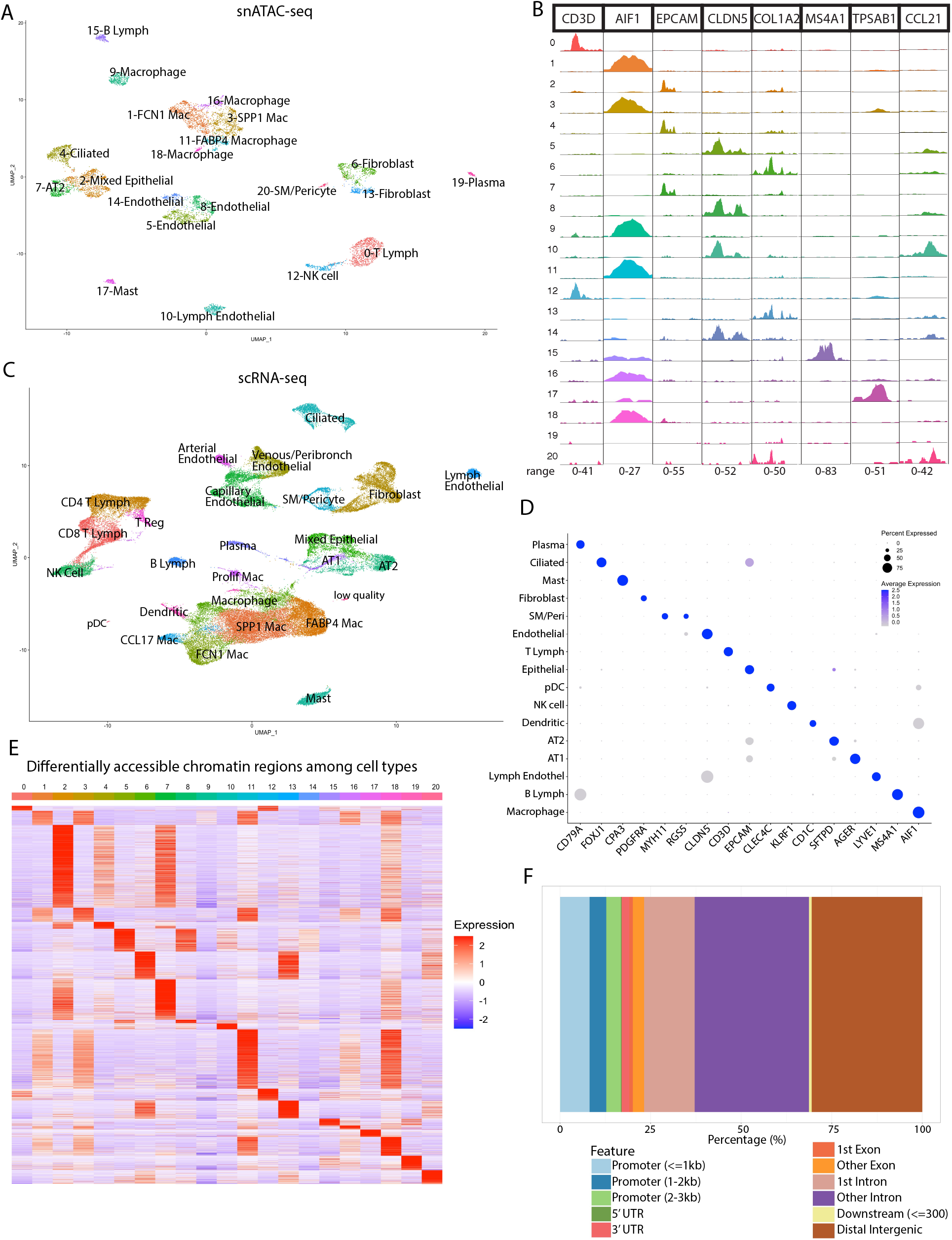
ScRNA-seq and snATAC-seq profiling of the human IPF and healthy lung. A) UMAP plot snATAC-seq dataset (n=3 IPF, 2 control) identified by cluster number and cell type. B) Merged coverage plots demonstrating pseudo-bulk chromatin accessibility (fragment coverage by frequency of Tn5 insertion) around marker gene promoters. Y-axis cluster numbers correspond to cell clusters in Figure 1A. Range of normalized accessibility for fragment coverage of each gene listed on x axis. C) UMAP plot of scRNA-seq dataset (n=10 IPF, 8 control) identified by cell type. D) Dot plot of scRNA-seq dataset showing gene expression of selected cell-type specific marker genes. Diameter of the dot corresponds to the proportion of the cells expressing the gene, and color density of the dot corresponds to the average expression level relative to all cell types. E) Heatmap of average number of Tn5 cut sites within the differentially accessible regions (each row is a unique DAR) for each cell type. Color scale represents a z-score of the number of Tn5 sites within each DAR. F) Bar plot of annotated DAR locations for each cell type.

We detected all major cell types within the lung with both the scRNA-seq and snATAC-seq datasets, with 260,166 accessible chromatin regions among 8,738 nuclei in the snATAC-seq dataset. In snATAC-seq, cell types can be distinguished by whether differentially accessible regions (DARs) of the chromatin are conformationally “open” or “closed.” To investigate alterations in chromatin accessibility between cell types, DARs with a log2-fold-change threshold of 0.25 occurring in at least 5% of nuclei within a cluster were identified, with epithelial and macrophage clusters having the most unique DARs (Figure 1E). Sequenced peak regions were annotated to the nearest gene and region of the genome with the R package ChIPSeeker (Figure 1F), with the majority of peaks in distal intergenic or intronic regions (16). The distribution of peak genomic regions was similar across cell types.

### Mesenchymal transcriptional and chromatin accessibility profiling

To more closely examine the myofibroblasts, we subclustered scRNA-seq and snATAC-seq fibroblast, smooth muscle, and pericyte populations to improve identification of the fibroblast subpopulations. By transcriptomes, we identified 3 major populations consisting of myofibroblasts, alveolar fibroblasts, and adventitial fibroblasts, and a minor population we refer to as *CXCL2*^hi^ fibroblasts (Figure 2A). Myofibroblasts originated primarily from IPF samples, while the non-myogenic fibroblast populations were observed in both IPF and donor control samples (Figure 2B). The myofibroblasts were defined by upregulation of *CTHRC1, POSTN, COMP,* and *COL3A1*, the alveolar fibroblasts by *SPINT2, FGFR4, GPC3,* and *MACF1* (analogous to our previously described *SPINT2*^hi^ fibroblasts), and the adventitial fibroblasts by upregulation of *PI16, MFAP5, IGFBP6* (analogous to our previously described *MFAP5*^hi^ fibroblasts) (Supp Fig 4A,5A)(17–19). In the snATAC-seq mesenchymal subclustering, 2 clusters of myofibroblasts, 3 clusters of non-myogenic fibroblasts, and a cluster of pericytes and smooth muscle cells were present (Figure 2C). An additional cluster contained nuclei with significant gene activity for both fibroblast and myeloid lineage markers, which were considered to be likely doublet nuclei and excluded from further downstream analyses. Control fibroblasts clustered distinctly from the IPF fibroblasts, which were divided into myofibroblast, alveolar, and adventitial fibroblasts (Supp Fig 4C). We focused our analysis to the comparison of IPF myofibroblasts to IPF non-myogenic fibroblasts (including the adventitial, alveolar, and *CXCL2*^hi^ fibroblast populations), as well as IPF fibroblasts to control fibroblasts. There were 163 DARs more accessible in IPF myofibroblasts vs 88 DARs more accessible in IPF non-myogenic fibroblasts (by Bonferroni adjusted p-value <0.05). Similarly, 19 DARs were more accessible in IPF fibroblasts vs 25 DARS in control fibroblasts. These DARs most commonly occurred in introns. Differentially accessible regions were overall consistent across individual IPF samples (Figure 2D). Of the 163 DARs more accessible in IPF myofibroblasts 17 were annotated to genes differentially expressed when comparing IPF myofibroblast to IPF non-myogenic fibroblasts, including *SPON2*, *PLEKHG1*, *SMYD3, AKAP7, LOXL2*, and *TSPAN2* amongst others. In comparing IPF to control fibroblasts, 5 of the 19 significantly more accessible regions were annotated to genes upregulated in IPF fibroblasts including *FBXL7, SPON2, ATP10D,* and *RUNX1.* While traditionally annotated to the nearest gene (by least base pairs distance), *cis-*regulatory regions do not inevitably regulate the nearest gene but may instead associate with more distant genes via long-range interactions. Such long-range interactions may facilitate the associations between promoters and enhancers via three-dimensional chromatin looping under the regulation of transcription factors (20). DARs not located near differentially expressed genes may in fact regulate more distant genes via such long-range interactions.

**Figure 2.**
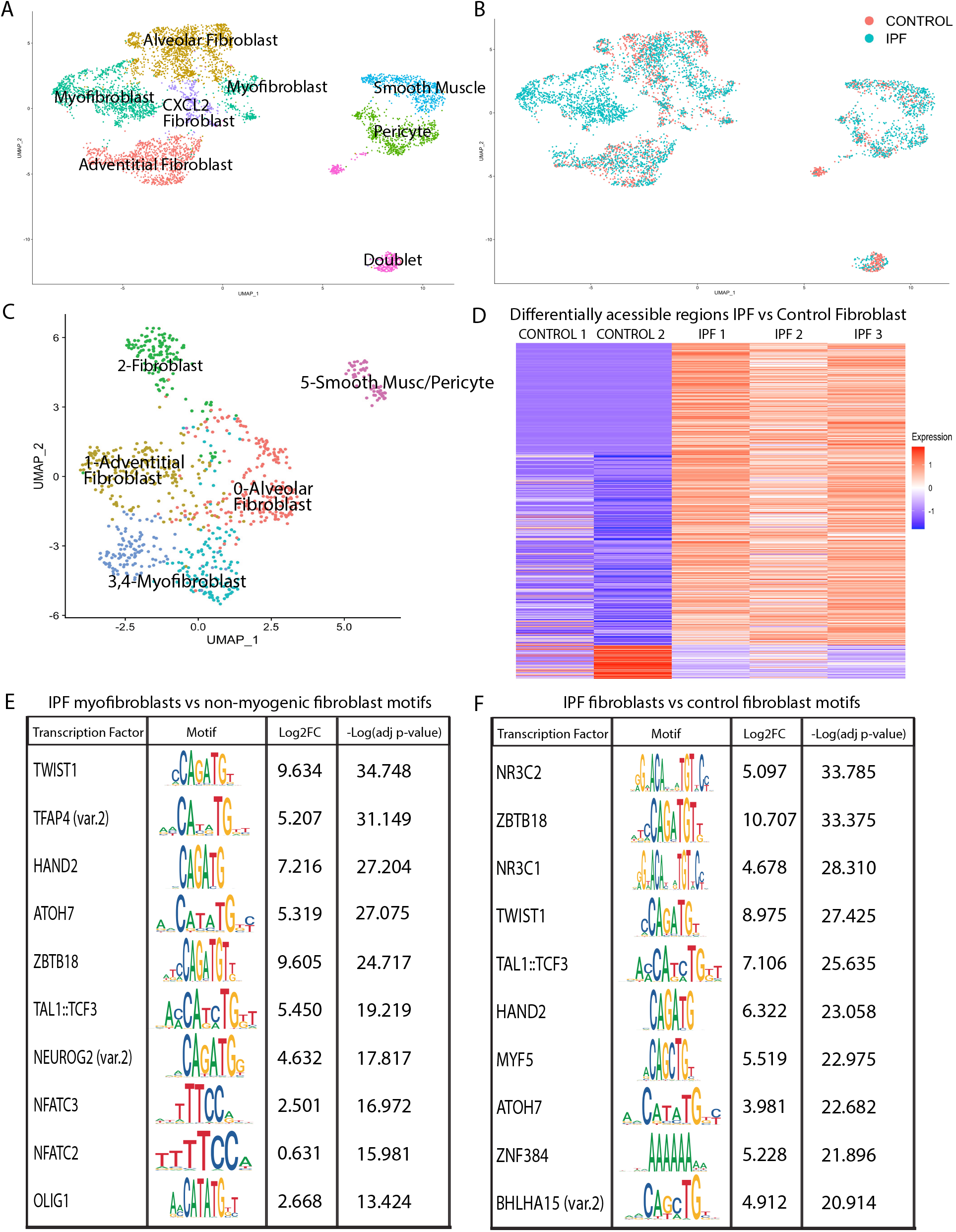
Fibroblast subpopulations and transcription factor motif activity in IPF myofibroblasts. A) UMAP plot of scRNA-seq fibroblasts, smooth muscle cells, and pericyte clusters by cell identity. B) UMAP plot of scRNA-seq fibroblasts, smooth muscle cells, and pericyte clusters from Figure 2A with cells depicted by origination from IPF vs control samples. C) UMAP plot of snATAC-seq fibroblast, smooth muscle cells, and pericyte clusters by cell identity. D) Heat map of average number of Tn5 cut sites within the differentially accessible regions when comparing IPF myofibroblasts to IPF non-myogenic fibroblasts, depicted by individual sample. Each row is a unique DAR. E) Transcription factors with the most significantly enriched motif activity when comparing IPF myofibroblasts to the IPF non-myogenic fibroblasts. F) Transcription factors with the most significantly enriched motif activity when comparing all IPF fibroblasts to all control fibroblasts.

### TWIST1 transcription factor motif activity enrichment in IPF myofibroblasts

TF motif enrichment can be inferred for groups of cells and cell types based on the enriched presence of TF-binding motifs within accessible chromatin regions, predicting critical active TFs regulating the cell state of interest. To assess TF-motif activity by cell population we used chromVAR to determine TF-associated accessibility in our snATAC-seq dataset. ChromVAR uses as input aligned fragment reads, chromatin-accessibility peaks, and chromatin motif position weight matrices and genomic annotations (21), and determines for each nuclei a bias-corrected “deviation” in motif accessibility (referred to as motif activity) from the anticipated motif accessibility based on the average of all nuclei. TF motif activity was defined for all nuclei, with differential TF motif activity examined between clusters, IPF myofibroblasts and non-myogenic fibroblasts, and IPF and control fibroblasts. Specific TF motifs were associated with each individual cell type. Known cell-type enriched TFs validated our data and analysis, such as FOXA1 and TEAD1 in alveolar epithelial cells (22), ETS1 in natural killer cells and T lymphocytes (23), and MEF2C in smooth muscle cells (24). Motifs highly enriched in fibroblasts compared to other cell types included ZBTB26, NFATC2, TWIST1, MYF5, HSF1, HSF2, and PBX2, amongst others (Supp Table 3).

On a global scale, TF expression correlated only to a limited degree with TF activity, supporting the notion that TFs may act as activators or repressors of gene expression based on post-transcriptional regulation of their activity. Of the 208 significant unique TF motif activities identified when comparing IPF to control fibroblasts, only 36 showed differential gene expression comparing these two cell populations. Comparing IPF myofibroblasts to non-myogenic fibroblasts, motifs for TWIST1, TFAP4, HAND2, ATOH7, and ZBTB18 were the most significantly enriched (Figure 2E), with *TWIST1, TCF3,* and *NFATC3* showing increased expression in myofibroblasts. Comparing IPF to control fibroblasts, motifs for NR3C2, ZBTB18, NR3C1, TWIST1, and TAL1::TCF3 were the most significantly enriched, with *NR3C1* conversely showing decreased expression in IPF fibroblasts (Figure 2F). Multiple TFs with a shared consensus binding sequence may bind to a highly similar motif, distinguished by only minor differences in their motif position weight matrices, such as the similar E-box motif of *TWIST1* and *HAND2.* ChromVAR and other *in silico* motif enrichment analyses alone cannot definitively determine which TF binds to a particular motif. As TWIST1 motif activity was significantly enriched in IPF myofibroblasts (Log2FC= 9.634, adjusted p-value=1.79E-35) and IPF fibroblasts (Log2FC=8.975, adjusted p-value=3.76 E-28) and its gene expression is highly specific to myofibroblasts (Figure 3A–3C), we further investigated TWIST1. Two regions of *TWIST1 (*within the promoter and a gap region), had statistically significant differential accessibility in comparing myofibroblast to non-myogenic fibroblasts (Figure 3D). *TWIST1* expression was upregulated in IPF myofibroblasts (Log2FC=3.136, adjusted p-value= 1.41E-24) vs non-myogenic fibroblasts.

**Figure 3.**
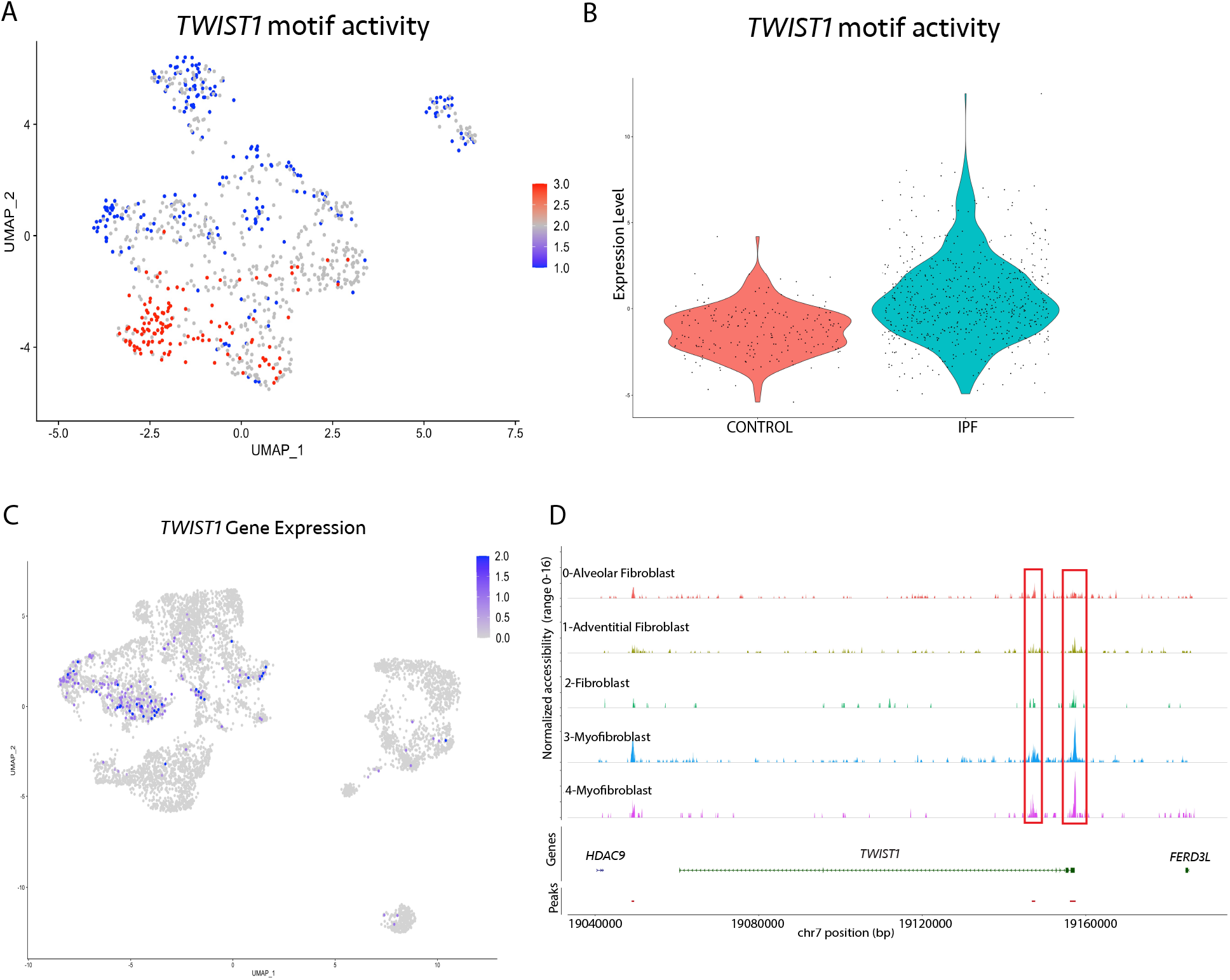
*TWIST1* expression and motif activity in IPF and control fibroblasts. A) UMAP plot of *TWIST1* motif activity in the mesenchymal snATAC-seq clustering, depicted by scaled expression with red representing the highest motif activity. B) Violin plot of *TWIST1* motif activity comparing IPF vs control fibroblasts only. C) UMAP plot of *TWIST1* gene expression in the mesenchymal scRNA-seq clustering demonstrating high *TWIST1* expression in the myofibroblasts only. D) Coverage plot demonstrating Tn5 insertion frequency by snATAC-seq fibroblast cluster in the *TWIST1* gene region. Red boxes represent regions of statistically significant differential accessibility in IPF myofibroblasts vs non-myogenic fibroblasts.

### Increased expression of Twist1 in collagen-producing cells is associated with increased collagen synthetic activity

We have previously observed that IPF patients with the highest expression of *TWIST1* by whole lung microarray analysis exhibited the most impaired gas exchange. Combining these data with our ATAC-seq observations above led us to consider how *Twist1* overexpression in the fibroblast compartment may impact an animal model of pulmonary fibrosis. We examined the effect of induced expression of Twist1 in lung *Col1a2*+ expressing fibroblasts (Twist1-LUC, Figure 4A) (25, 26). Lung fibroblasts were isolated and cultured in the presence of tamoxifen and doxycycline, to induce Twist1 expression, with and without TGFβ. In unstimulated Twist1-LUC fibroblasts, we observed increased expression of collagen I and the myofibroblast marker α-smooth muscle actin (α-SMA) compared to wild type fibroblasts (Twist1-WT) (Figure 4B). This effect was amplified in the presence of TGFβ. These data indicate that Twist1 expression in lung fibroblasts is associated with increased expression of collagen I and α-SMA. Next, we explored the effect of increased Twist1 in fibroblasts following bleomycin injury (Figure 4C, D). Following bleomycin, we grossly observed a comparable degree of acute lung injury in both Twist1-WT and Twist1-LUC. We incidentally noted the presence of airspace multinucleated giant cells in Twist1-LUC mice. When we quantified acid-soluble (fresh) collagen synthesis, we observed increased collagen in bleomycin-injured Twist1-LUC mice compared to Twist1-WT (Figure 4D). Taken together, these data suggest that increased expression of Twist1 in collagen-producing cells is associated with increased collagen synthetic activity in both *in vitro* and *in vivo* models.

**Figure 4.**
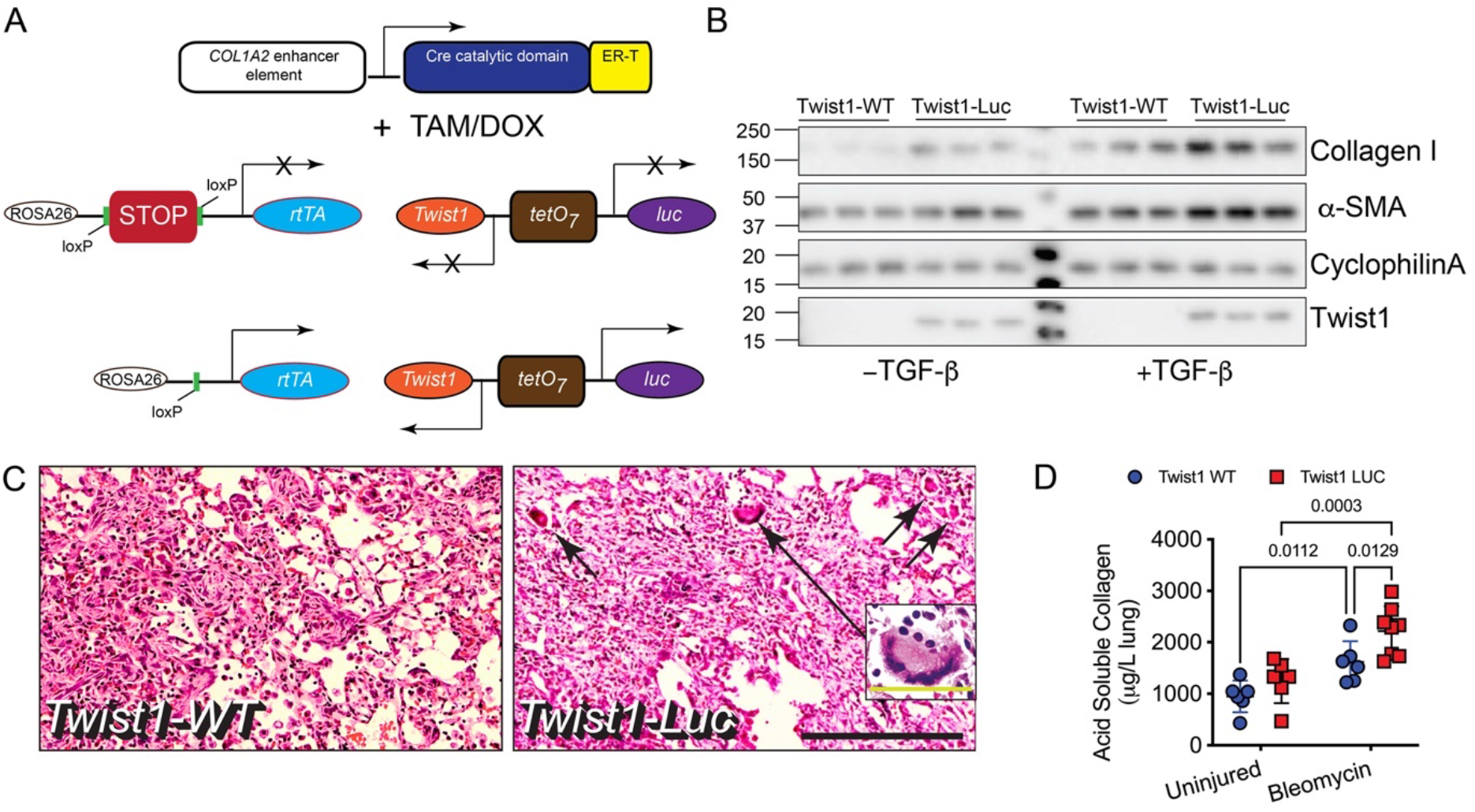
Overexpression of Twist1 in *Col1a2+* cells leads to increased collagen I levels *in vitro* and *in vivo.* A) A triple transgenic animal was bred where Cre recombinase is under control of the *col1a2* enhancer element (*col1a2-Cre-ER(T)*). In the presence of tamoxifen (TAM), the STOP signal is excised leading to expression of the reverse tetracycline transactivator (rtTA). In the presence of doxycycline and the rtTA, the tetO7 operator is activated leading to expression of Twist1 and luciferase. B) Lung fibroblasts from Twist1-WT (wild type) and Twist1-Luc (Twist1 overexpressors) were incubated in the presence of TAM and DOX with and without TGFβ (2ng/mL). Cells were lysed and subjected to immunoblotting for Collagen I, α-smooth muscle actin (α-SMA), Twist1, and the loading control, cyclophilin A. In the presence of TAM/DOX, increased collagen, α-SMA, and Twist1 in Twist1-LUC fibroblasts. This was amplified in the presence of TGFβ (N=3). C) Twist1-WT and Twist1-Luc mice were injured with bleomycin. Animals were sacrificed at 14d. A comparable degree of histologic injury was observed in Twist1-WT and Twist1-Luc mice (x100 magnification, inset bar = 200μm). Incidentally noted multinucleated giant cells (MGC) are identified by the black arrows. The long black arrow magnifies one MGC (x400 magnification, inset yellow bar=50μm). D) Determination of acid-soluble collagen content showed a significant increase in bleomycin-induced collagen in Twist1-Luc mice compared to Twist1-WT (by two-way ANOVA, with Fisher’s LSD post-hoc test, N=6-8).

### RNA-seq analysis of mouse lung fibroblasts with *Twist1* overexpression is associated with dysregulation of multiple fibrosis-relevant genes

To further explore genes regulated by Twist1 *in vitro*, we compared gene expression in cultured lung fibroblasts from three Twist1-LUC mice and three Twist1-WT mice *in vitro* by RNA sequencing (Figure 5). A heatmap was generated and identified a significant number of genes that were differentially expressed between Twist1-LUC and Twist1-WT fibroblasts. One of three of Twist1-WT fibroblasts clustered more closely to Twist1-LUC fibroblasts. By immunoblotting, this line had significantly higher Twist1 protein expression than the other two Twist1-WT lines but less than the Twist1-LUC fibroblasts, representing an intermediate phenotype. Volcano plot analysis (Figure 5B-D) identified several significantly up-regulated genes that have been associated with TGFβ signaling and pulmonary fibrosis including *ltbp1* (27)*, tbx4* (28)*, tnc* (29, 30) *and thbs4* (31). Dysregulated canonical pathways including “Systemic Lupus Erythematosus in B cell Signaling” (*tnfsf11*, and *tnfsf15*) and “Role of Hypercytokinemia in the Pathogenesis of Influenza” as well as HIPPO signaling—which has been implicated in pulmonary fibrosis (32)— and pulmonary and hepatic fibrosis pathways (*ptch2*, *il1rap*, *itgb3*, and *flt3*). Downregulated signaling for xenobiotic metabolism, glutathione-mediated detoxification and NRF2-mediated oxidative stress response (*gsta3* (33)*, nqo1* (34*), cyp1a1* (35), and *acta1* (36)) is supportive of previous data suggesting that loss of these pathways is a component of fibrosis (37, 38). When a heat map was generated based on the genes that showed the highest coefficient of variation, fibrotic genes such as *col1a1*, *col1a2*, *col3a1*, *tnc*, *thbs1*, and *lox* were highly upregulated in Twist1-LUC fibroblasts (Supp Fig 7). When we compared our snATAC-seq data above with our RNA-seq analysis of mouse lung fibroblasts, we observed that *MMP8* and *TNFRSF9,* which were *a*mongst the top upregulated genes in Twist1-LUC fibroblasts, have increased chromatin accessibility in IPF myofibroblasts (Supp Fig 8) supporting the observation that downstream targets of TWIST1 have altered chromatin in myofibroblasts.

**Figure 5.**
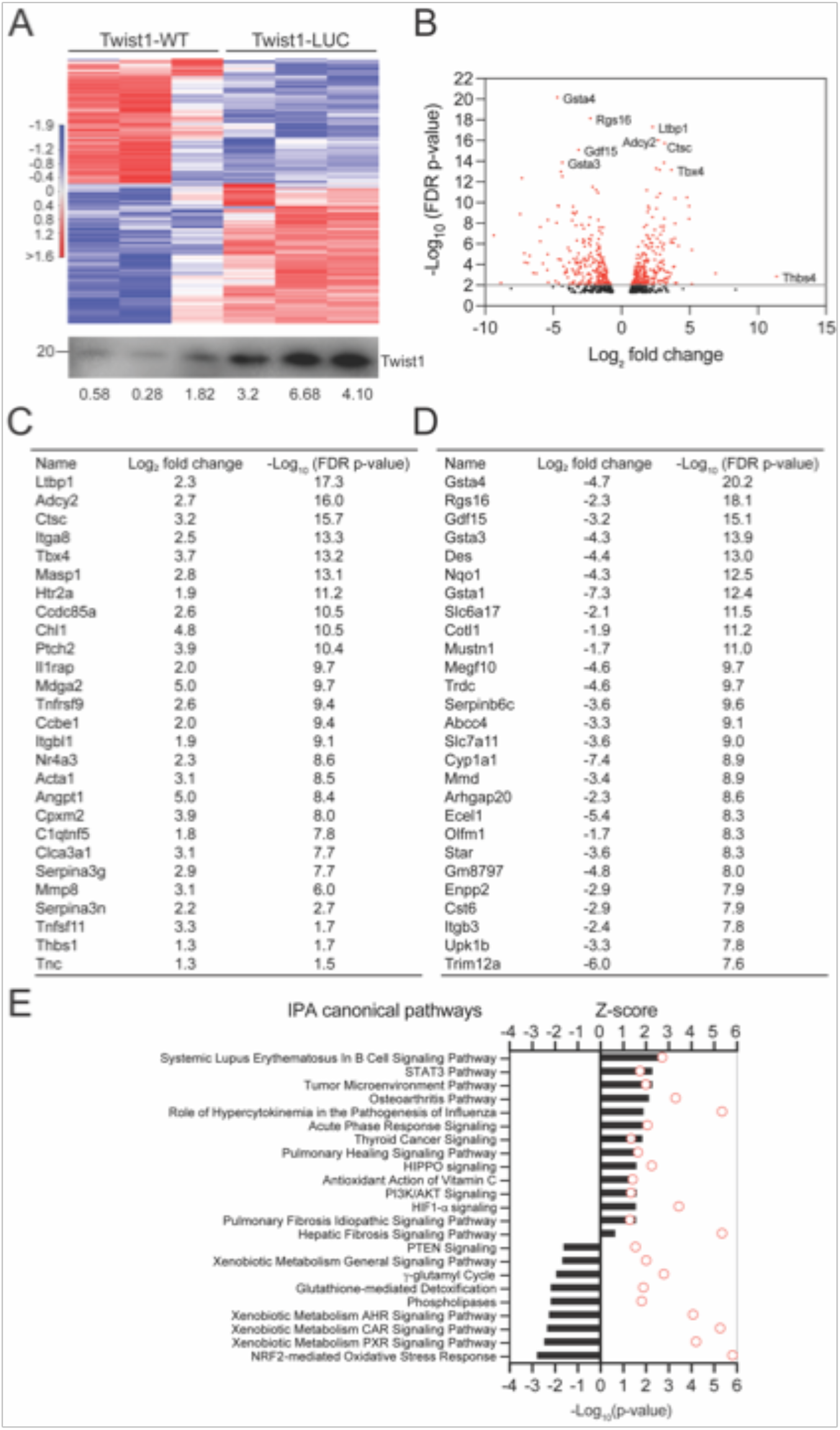
Twist1 overexpression in mouse lung fibroblasts is associated with dysregulation of several pulmonary fibrosis genes and pathways. Bulk RNA-seq was performed on fibroblasts isolated from lungs of WT and Twist1-LUC mice (N=3). Estimation of differential gene expression using CLC genomics workbench was performed comparing fibroblasts from Knock-in with normal lungs. A) Hierarchical clustering heatmap of significant differentially expressed genes was generated using CLC genomics workbench using minimum absolute fold change of 3.0 and FDR p-value threshold of 0.05. Immunoblotting is shown for the individual lines subjected to RNA-seq. Densitometry normalized to β-actin is shown beneath. B) Volcano plot shows comparative analysis of differentially expressed genes between the WT and Twist1-LUC. C) List of Dysregulated genes by FDR that are up-regulated ranked by - Log10 (False discovery rate p-value) with p<0.05 cut-off. D) List of top downregulated genes ranked by −Log10 (False discovery rate p-value) with p<0.05 cut-off down-regulated in Twist1-LUC fibroblasts compared to Twist1-WT fibroblasts. E) Ingenuity pathway analysis of dysregulated canonical pathways between WT and Twist1-LUC (n=3) by z-score and FDR.

## Discussion

Our study demonstrates that the differentiation of myofibroblasts—the central effector cells in IPF—is characterized by a significant shift in chromatin accessibility dominated by opening of E-box TF binding sites. We utilized single-cell sequencing platforms of *ex vivo* IPF lungs to obviate the distortion of signals across heterogeneous populations and the changes in chromatin accessibility and gene expression that may occur with expansion in culture. And of the E-box TF binding sites, we identified TWIST1 as the most highly enriched regulator of myofibroblast activity. We then confirmed a critical regulatory role for TWIST1 by demonstrating that overexpression of Twist1 in the fibroblast compartment, *in vitro* and *in vivo*, led to increased expression of collagen I and α-SMA.

Previous studies have identified epigenetic changes in IPF lungs via methylation profiling (39–41), however knowledge of cell type specific epigenetic alterations remained limited. A recent study by Hanmandlu *et al.* utilized bulk ATAC-seq of cultured fibroblasts to investigate chromatin accessibility in IPF upper lobe fibroblasts (42). They similarly identified the significant enrichment of the E-box TFs TWIST1 and ZBTB18 binding motifs in IPF compared to control fibroblasts, further supporting an important role for TWIST1 and the E-box TFs in IPF myofibroblasts. However, other TF motifs implicated by their bulk ATAC-seq analysis including FOXA1 and FOXP1 were significantly less enriched in our analyses, while CBFB was enriched in IPF fibroblasts in the bulk analyses only, indicating accessibility for these TF motifs may be altered by *in vitro* culture, though heterogeneity amongst clinical samples cannot be ruled out. The E-box TF MYF5, a known regulatory factor critical to myogenic differentiation during embryogenesis, was also enriched in our snATAC-seq, but not the prior cultured fibroblasts.

The consistent enrichment of E-box TF motifs in IPF myofibroblasts, suggests modulation of their accessibility as a critical step within or resulting from activation of the aberrant myofibroblast program. TWIST1 and other basic Helix Loop Helix (bHLH) TFs bind to canonical E-box sequences in order to regulate gene expression. Their bHLH domain permits dimerization with other bHLH TFs, altering the transactivating function of such partners (43). While we focus in this work on confirmation of a role for TWIST1, the complex mechanism of myofibroblast activation undoubtedly involves the coordinated activation of multiple TFs. Defining such shifts in a cell population’s epigenetic state now opens the door for novel molecular and computational approaches to therapeutic development in IPF. For instance, the rapid advancement of small molecule therapeutics including those inhibiting DNA-protein binding and TF complexes (44) supports identifying and targeting key bHLH dimerization pairs or other coordinated TF binding partners of TWIST1. Investigating chromatin accessibility changes in the context of drug therapies also has the potential to identify agents halting (and ideally reversing) such epigenetic changes, potentially as an early signal for therapeutic response in the context of early phase clinical trials. In contrast to previous studies (45–48) that have examined the role of TWIST1 in fibroblasts in cancer and fibrosis, to our knowledge, we are the first to report that fibroblast-specific overexpression of Twist1 *in vivo* is associated with increased collagen synthetic activity. This corroborates our observation that increasing gene expression of *TWIST1* in IPF is associated with more compromised gas exchange (46). It was striking that one Twist1-WT line of fibroblasts, through experimental variation, mapped more closely with the Twist1-LUC fibroblasts, suggesting a very narrow dynamic range of expression of Twist1. Deviations from that narrow range of expression can lead to pronounced differences in fibroblast phenotypes (46). Taken together, these data support a unique model whereby *TWIST1* expression serves as a critical molecular “rheostat” in IPF. Cellular levels of TWIST1 are thus tightly regulated in cells and even small changes in Twist1 levels can significantly impact the fibrotic phenotype. Prior murine studies have implicated YAP1/TEAD1 regulation of Twist1 expression as an important upstream mechanism of Twist1’s fibrotic effects (49), however the distinct enrichment of TWIST1 binding sites in IPF myofibroblasts (rather than those of YAP1 or TEAD1) along with its upregulated expression suggest it may be a critical juncture for altering fibrosis.

Surprisingly, we observed multinucleated giant cells (MGC) in bleomycin-injured Twist1-LUC mice. This curious observation is not a common finding in bleomycin injury and has not been previously described. Consistent with the capacity of fibroblasts to modulate the inflammatory response *in vivo*, we did observe significantly increased expression of Receptor activator of NF-κB Ligand (RANKL, encoded by *TNFSF11*) signaling—which is essential for MGC formation in bone (50)—in mouse lung fibroblasts, *in vitro*, following overexpression of Twist1. While the presence of MGC is not a feature of IPF, it is a feature of other fibrotic lung diseases such as hypersensitivity pneumonitis (51) and hard metal pneumoconiosis (52). Thus, TWIST1 may play a particularly important role in these fibrotic lung diseases, warranting future study.

As the demonstrated changes in *TWIST1* motif enrichment were consistently observed despite the modest sample size, it merited further mechanistic investigation. Despite small snATAC-seq sample numbers, we demonstrate consistency of differentially accessible regions between cell types and IPF and control fibroblasts across individual samples. Future expansion to larger snATAC-seq datasets may allow enhanced assessment of covariance between accessible chromatin sites to predict *cis*-regulatory interactions and further link regulatory regions to their target genes (53). All IPF samples were from patients with end-stage disease receiving care at a tertiary medical center and may not reflect the comprehensive IPF population. Due to the limited sample number, we were unable to perform covariate analyses for age or sex. To limit confounding from differences in experimental technique, we have chosen to analyze only snATAC-seq samples processed using the same platform and reagent chemistry.

In summary, our analysis utilizes human multiomic single-cell analyses combined with *in vivo* murine disease models to investigate TF gene regulatory networks critical to myofibroblasts in IPF. Comparison of *in vivo* IPF myofibroblasts to non-myogenic and control fibroblasts identified a dynamic opening of E-box TF binding sites in the activated IPF myofibroblasts, with the E-box TF TWIST1 particularly implicated as a positive regulator of myofibroblast activity. Both low (46) and high expression of Twist1 in fibroblasts is associated with increased collagen deposition in the lung, confirming its role as a critical regulator in the fibrotic lung. Future studies delineating the global mechanism modulating E-box TF motif accessibility may identify crucial therapeutic targets for deactivating the aberrant myofibroblast program.

## Methods

### Materials

All scRNA-seq and snATAC-seq reagents were purchased from 10X Genomics (Pleasanton, CA). APC-conjugated antibodies to CD45 and CD326 were purchased from Biolegend (San Diego, CA). Anti-APC Microbeads were purchased from Miltenyi (Bergisch Gladbach, Germany). Monoclonal antibody against Twist (clone 2C1a) was purchased from Abcam (Cambridge, MA). Antibodies to Collagen type I were obtained from Rockland (Limerick, PA). An antibody against cyclophilin A was purchased from Santa Cruz Biotechnology (Dallas, TX). Sircol collagen assay kit was purchased from Biocolor (Belfast, UK).

### ScRNA-seq/snATAC-seq

The University of Pittsburgh Institutional Review Board approved procedures involving human samples. Explanted subpleural peripheral lung tissue was digested to single-cell suspensions using Liberase DL (Roche) and bovine pancreas DNAse I (Sigma-Aldrich) as previously described (17). Three samples were partially enriched for fibroblasts using magnetic-activated depletion of CD45 (PTPRC) and CD326 (EPCAM) positive cells, with 50/50 loading of the enriched and non-enriched cell suspension at the time of running scRNA/snATAC-seq. Single-cell suspensions were split, with a portion used for performing scRNA-seq as previously described (19), and the remaining suspension used for nuclei generation. The 10X Genomics low cell input nuclei isolation protocol was followed with alteration of lysis timing as follows. To isolate nuclei, a cell suspension of 50,000 cells was centrifuged at 300 rcf for 5 minutes at 4°C, followed by resuspension of the cells in chilled PBS containing 0.04% BSA for 1,000 cells/*μ*l cell suspension. The single cell suspension was centrifuged again at the same settings, followed by removal of supernatant. Cells were mixed with 45 *μ*l of chilled lysis buffer and incubated for 90 seconds on ice, followed by addition of 50 *μ*l of of chilled wash buffer. Nuclei suspension was centrifuged at 500 rcf for 5 minutes at 4°C, washed with diluted nuclei buffer and centrifuged again. Nuclei pellet was resuspended in diluted nuclei buffer for cell counting and performing snATAC-sequencing. 2000 nuclei of each lung sample were loaded for transposition with Chromium single-cell ATAC system “E” chips (10X Genomics). Barcoded libraries were pooled and sequenced on an Illumina NovaSeq 6000 at the UPMC Genome Center. DNA libraries were sequenced to a depth of 400M read pairs per library.

CellRanger ATAC pipeline (v1.2.0; 10X Genomics) with the hg19 human genome reference was used for initial read processing, library quality control, peak calling and cell calling for individual samples. The number of nuclei recovered was 1,776-2,832 per library with the median unique fragments per nucleus 8,018-20,487. The R package Signac (v1.3.0) (15) was used for merging samples, unsupervised clustering, and downstream analyses. A unified peak set was created and quantified in each dataset, followed by new fragment objects, Seurat objects, and merging of the 5 samples. Batch correction at the individual sample level was performed using the R package harmony (v1.0) (13). Peak calling was performed by cluster using macs2 (54), with this peak set used for downstream analysis and subclustering of the mesenchymal populations. For manual annotation of principal cell types, the UMAP projection of nuclei was queried for clusters showing accessibility of known cell type-specific genes, and further verified by performing cell type predicted labeling using the scRNA-seq dataset as the annotation reference, at both the all cell types and mesenchymal subclustering level. Differentially accessible regions between clusters and other specified groupings were calculated by logistic regression test with number of peak region fragments as the latent variable and requiring peak accessibility in 5 percent or more of nuclei in at least one population. ‘Gene accessibility’ for the nuclei was calculated by aggregating total fragments from the gene body and 2kb upstream of the transcription start site. Signac was used for display of pseudo-bulk coverage tracks of Tn5 integration for designated cell populations.

Differential motif accessibility was performed using the R package chromVAR (v1.12.0) (55) with motifs from the cisBP database. In brief, ‘motif activity’ is calculated for each motif within each nuclei by comparing motif occurrence in accessible peaks to motif occurrence in a set of background peaks matched for GC content. To calculate differential motif accessibility between populations a logistic regression test was used with number of peak region fragments and macs2 peak fragments as latent variables.

For scRNA-seq analysis CellRanger pipeline with the GRCh38 human reference genome was used. The R package Seurat (v4.0.3) was used for downstream filtering, integration, unsupervised clustering, and analysis. Cells were filtered for greater than 200 genes, less than 50,000 UMIs, and less than 15 percent mitochondrial genes. Batch correction at the individual sample level was performed by reciprocal PCA integration when combining the 18 samples. Wilcoxon rank sum test with Bonferroni FDR correction was used for differential gene expression testing. SCTransform normalization followed by harmony batch correction was used for the mesenchymal subclustering. The raw data have been deposited in NCBI’s Gene Expression Omnibus.

### Twist1 conditional knockout mice

ROSA26-STOP tdTomato+ mice were purchased from Jackson (Bar Harbor, ME). Mice expressing cre recombinase in collagen-expressing cells (COL1A2) in the presence of tamoxifen were generously provided by Dr Benoit DeCrombrugghe (MD Anderson Cancer Center)(56). Doxycycline (DOX)-inducible mice that overexpress Twist1 have been described previously(25, 26). All mice were subjected to intraperitoneal (IP) injection of tamoxifen (80mg/kg for 5 days) at 5 weeks and 6 additional injections delivered three times per week beginning at 8 weeks of age during bleomycin injury experiments.

### Bleomycin-Induced Lung Injury

8-week-old mice were anesthetized with isoflurane in an anesthesia chamber. Bleomycin was administered at 2U/kg by transnasal aspiration (57). Mice were sacrificed on day 14 (57).

### Sircol Assay for Acid-Soluble Collagen

Mouse lungs were perfused through the right ventricle with PBS. The left lungs were excised and flash-frozen in liquid nitrogen, followed by lyophilization in preparation for collagen determination by the Sircol collagen assay. Lungs were homogenized in 0.5M acetic acid with protease inhibitors (Sigma-Aldrich). The homogenate was pelleted, and the supernatant was run across a QIAshredder column (Qiagen). The lung-acetic acid mixture was incubated with the Sircol Red reagent. The collagen-dye complex was pelleted, and the precipitated material was re-dissolved in NaOH. Optical density at 540 nm (OD540) was recorded using a microplate reader.

### Isolation of mouse lung fibroblasts and immunoblotting

Mouse lung fibroblasts were isolated from uninjured lungs of wild type and transgenic mice as described previously (57). Fibroblasts were lysed for immunoblotting to measure the protein expression of Twist1, Collagen I, α-SMA, and the housekeeping protein cyclophilin A. Immunoblotting was performed as previously described (58). Cells were treated with brefeldin A at one hour prior to lysis to inhibit secretion of collagen.

### Bulk RNA-seq analysis

Primary lung fibroblasts from Twist1-Luc, ColCre+, Rosa26RTta and their littermate controls (ColCre+, Rosa26RTta) were cultured (n=3) and further treated with Doxycycline and Tamoxifen. Total RNA was purified using RNAeasy kit (Qiagen) according to manufacturer’s protocol. RNA quality was determined using NanoDrop and Bioanalyzer. Stranded total RNA-seq libraries were prepared following ribosomal RNA depletion with Illumina reagents, and libraries were sequenced with an Illumina Nextseq500 sequencer with a depth of paired-end 20 million read pairs per sample at UPMC core (Pittsburgh, PA). Fastq files with high quality reads (phred score >30) were uploaded to the CLC Genomics Workbench (Qiagen). Reads were aligned to Mus musculus ensembl_v80 genome. Transcript counts, quality control, alignment, differential expression analysis, and preliminary enrichment analysis were performed using CLC genomics workbench. Volcano plot was generated using GraphPad Prism by plotting −Log10 (FDR p-value) on Y-axis and Log2 fold change on X-axis. Hierarchical clustering algorithm (heat map) was generated using CLC genomics workbench by using Euclidean distance and complete linkage as parameters and filtered by using the differential expression gene set with minimum absolute fold change of >3.0 and FDR p-valve threshold of p<0.05 in figure 5. The supplemental heatmap (Supp Fig 7) was generated using fixed number of features option where given number of features with the highest coefficient of variation (the ratio of the standard deviation to the mean) were kept. The raw data have been deposited in NCBI’s Gene Expression Omnibus. Additional enrichment analyses were conducted using Ingenuity Pathway Analysis (Qiagen).

## Study Approval

This work was approved by the Institutional Review Board and the Institutional Animal Care and Use Committee of the University of Pittsburgh.

## Author Contributions

EV designed research studies, conducted experiments, acquired data, analyzed data, and wrote the manuscript. HB conducted experiments, acquired data, analyzed data, and wrote the manuscript. JT designed research studies, conducted experiments, acquired data, analyzed data, and wrote the manuscript. TT conducted experiments, acquired data, and analyzed data. DIS analyzed data. JS conducted experiments and acquired data. LF acquired data and provided reagents. KC acquired data and provided reagents. MR acquired data and provided samples. AL acquired data and provided reagents. DWF provided hybrid mouse and wrote the manuscript. PTT provided hybrid mouse and wrote the manuscript. DJK designed research studies, acquired data, analyzed data, and wrote the manuscript. RL designed research studies, analyzed data and edited the manuscript. EV listed first of the equal contribution first-authors for primary compilation of the submission.

## Acknowledgements

Support for the studies was provided by R01 HL 126990 to DJK and P50 AR 060780-06A1 to RL and DJK. The authors would like to acknowledge the Center for Organ Recovery & Education (CORE) as well as organ donors and their families for the generous donation of tissues used in this study.

