## Supplemental Figures for "Single-Nucleus Chromatin Accessibility Identifies a Critical Role for TWIST1 in Idiopathic Pulmonary Fibrosis Myofibroblast Activity"

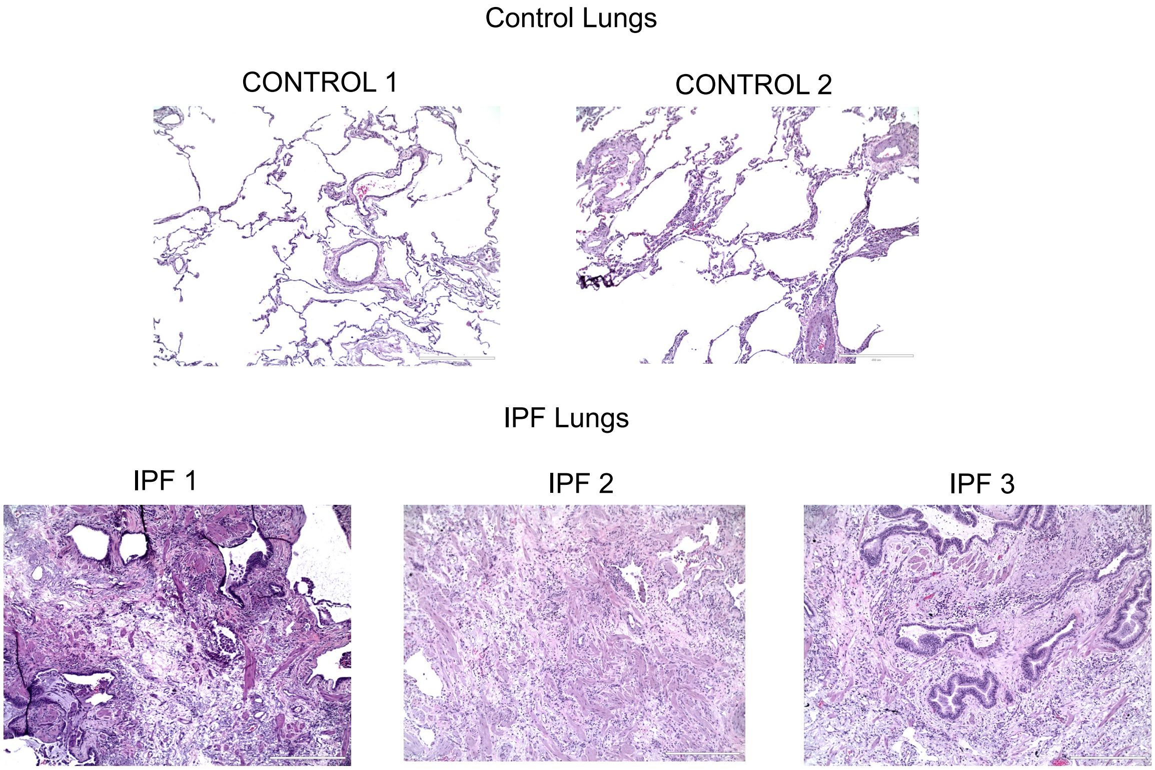


**Supplemental Figure 1. Histopathology of snATAC-seq explanted lung samples.**
H&E stained histopathology of adjacent lung tissue sections from the explanted lungs, identified by sample id. Scale bar=400$\mu$m. IPF, idiopathic pulmonary fibrosis.

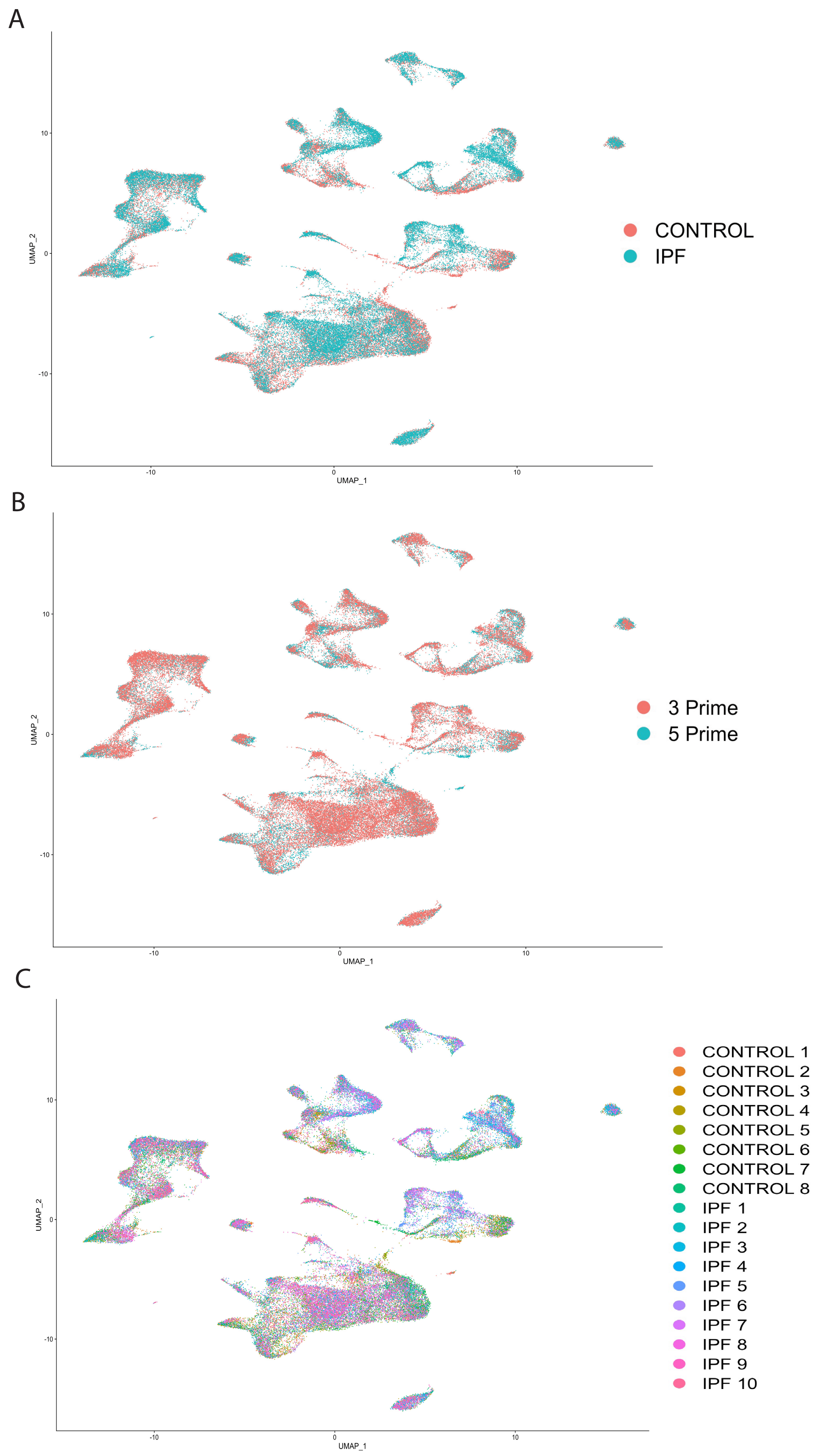


**Supplemental Figure 2. ScRNA-seq profiling of the human IPF and control lungs.** A) UMAP plot of scRNA-seq dataset (n=10 IPF, 8 control) depicted by origination from healthy control or IPF lungs. B) UMAP plot depicted by origination from 3’ or 5’ 10X Genomics chemistry. C) UMAP plot depicted by sample of origin.

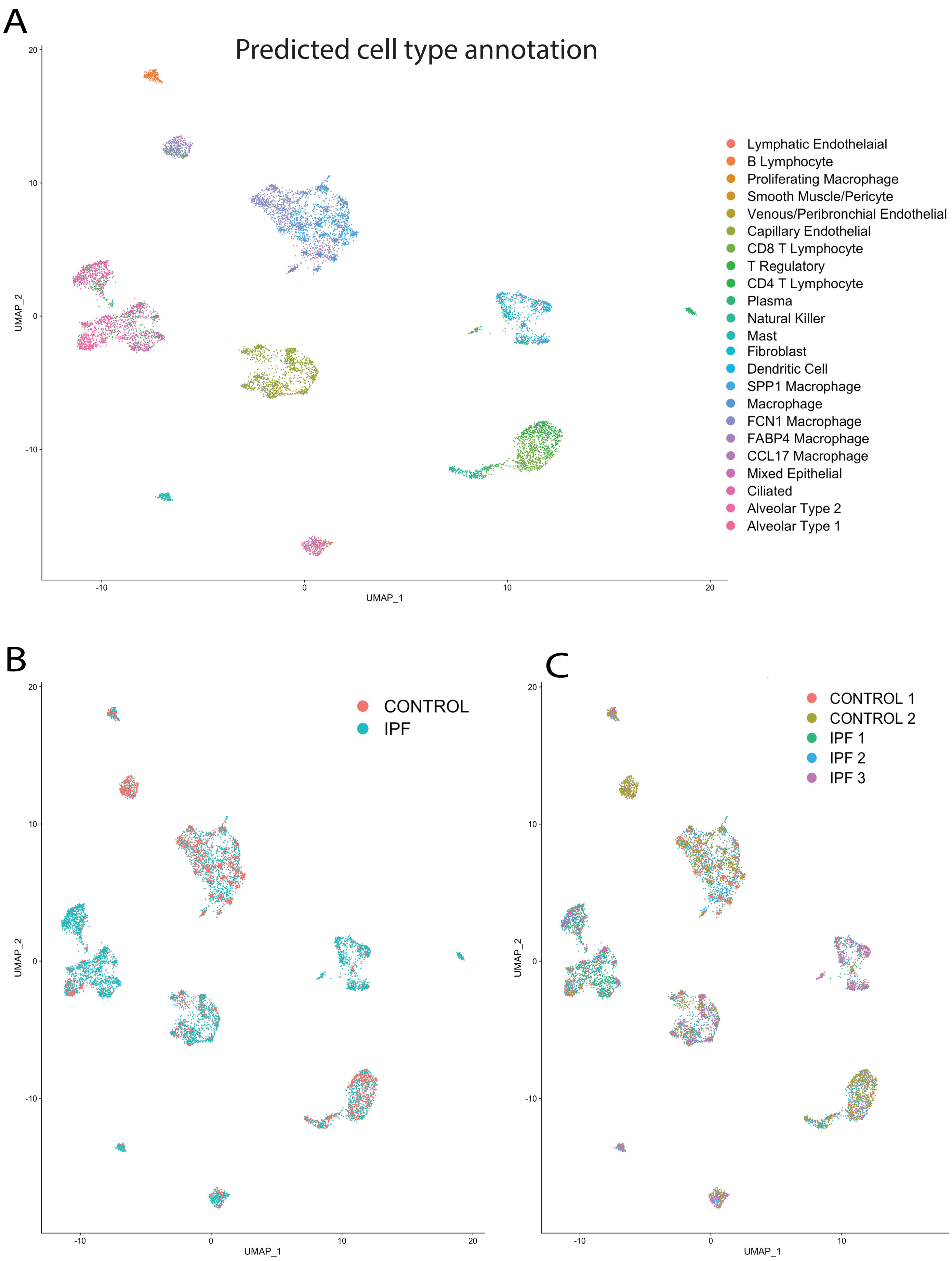
A
Predicted cell type annotation
Lymphatic Endothelaial
B Lymphocyte
Proliferating Macrophage
Smooth Muscle/Pericyte
Venous/Peribronchial Endothelial
Capillary Endothelial
CD8 T Lymphocyte
T Regulatory
CD4 T Lymphocyte
Plasma
Natural Killer
Mast
Fibroblast
Dendritic Cell
SPP1 Macrophage
Macrophage
FCN1 Macrophage
FABP4 Macrophage
CCL17 Macrophage
Mixed Epithelial
Ciliated
Alveolar Type 2
Alveolar Type 1
UMAP_1
UMAP_2
B
CONTROL
IPF
C
CONTROL 1
CONTROL 2
IPF 1
IPF 2
IPF 3

**Supplemental Figure 3. SnATAC-seq profiling of the human IPF and control lungs.**
A) UMAP plot of snATAC-seq dataset (n=3 IPF, 2 control) depicted by predicted cell-type annotation based on transfer of scRNA-seq dataset. B) UMAP plot depicted by origination from healthy control or IPF lungs. C) UMAP plot depicted by sample of origin.

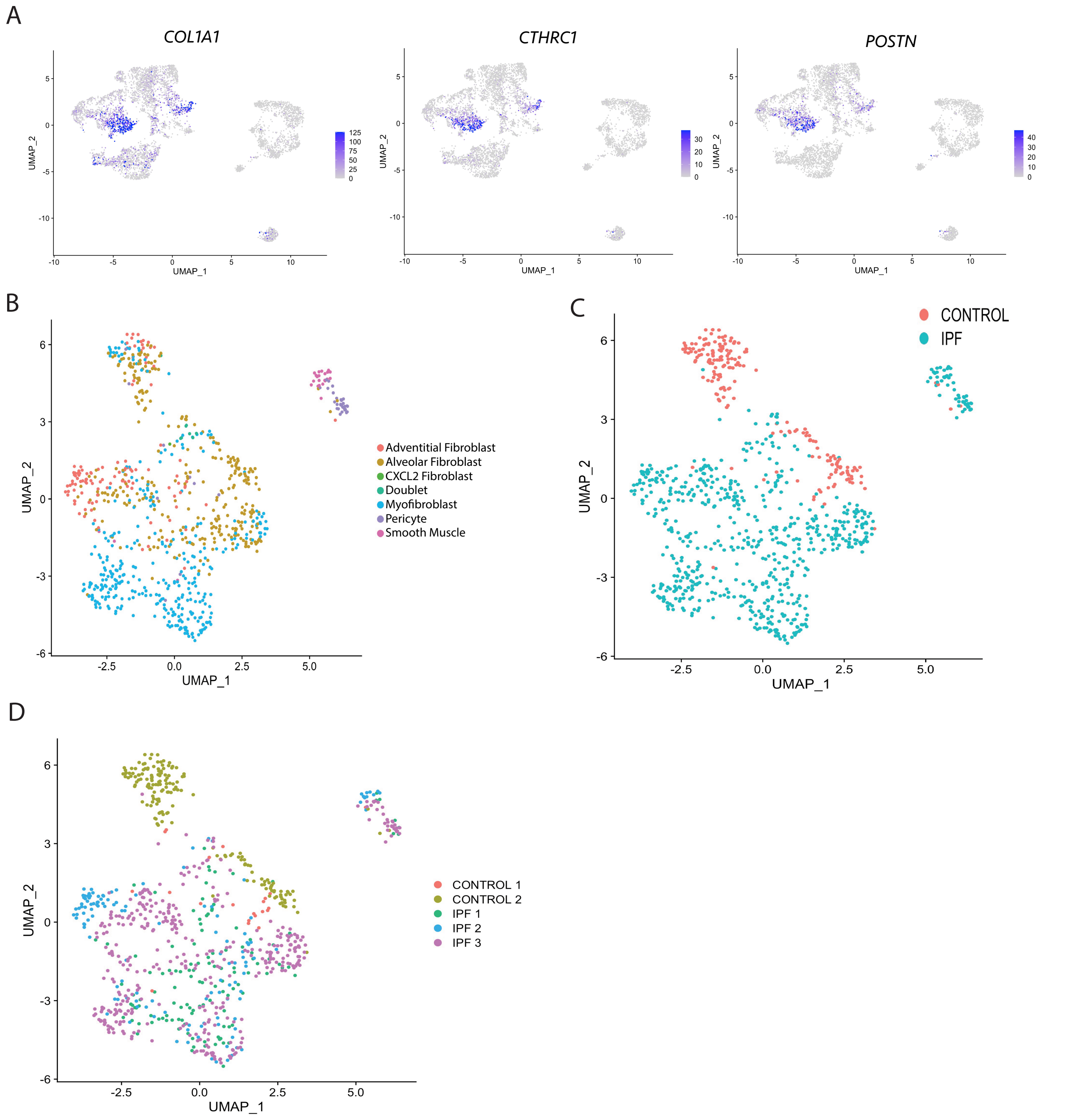


**Supplemental Figure 4. Mesenchymal cell subclustering.**

A) Feature plot of scRNA-seq mesenchymal subclustering with gene expression of *COL1A1, CTHRC1,* and *POSTN* depicting high expression in the myofibroblasts. B) UMAP plot of snATAC-seq mesenchymal subclustering depicting predicted cell-type annotation based on transfer of scRNA-seq dataset. C) UMAP plot of snATAC-seq mesenchymal subclustering depicted by origination from healthy control or IPF lungs. D)UMAP plot of snATAC-seq mesenchymal subclustering depicted by sample of origin.

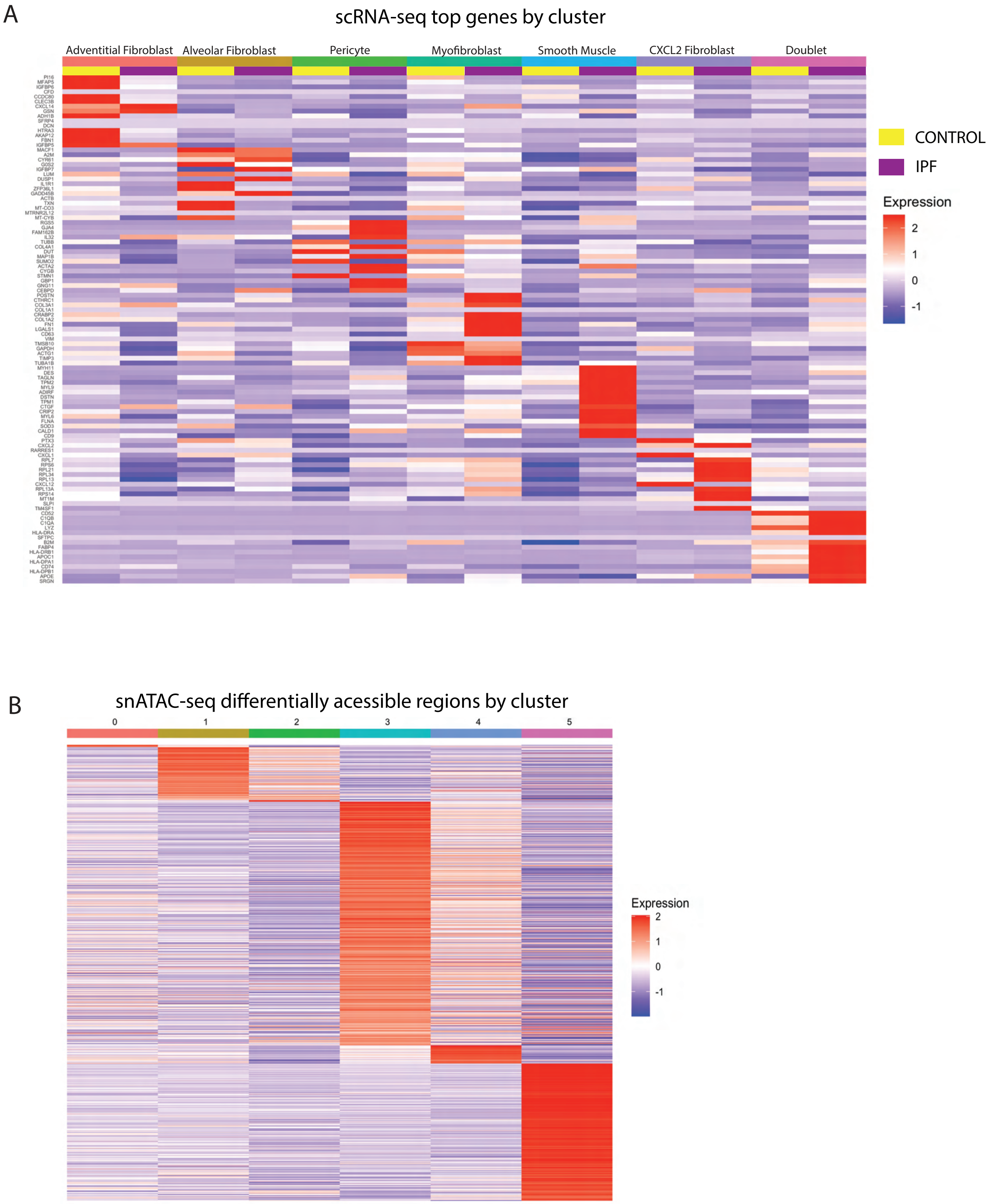


**Supplemental Figure 5. Mesenchymal cell subclustering**

A) Heatmap of average gene expression by cluster and disease status for top genes by mesenchymal cluster. B) Heatmap of average number of Tn5 cut sites within the differentially accessible regions (each row is a unique DAR) for each mesenchymal cluster. Color scale represents a z-score of the number of Tn5 sites within each DAR.

A

Top 50 Chromvar motifs by Cell type (average expression)

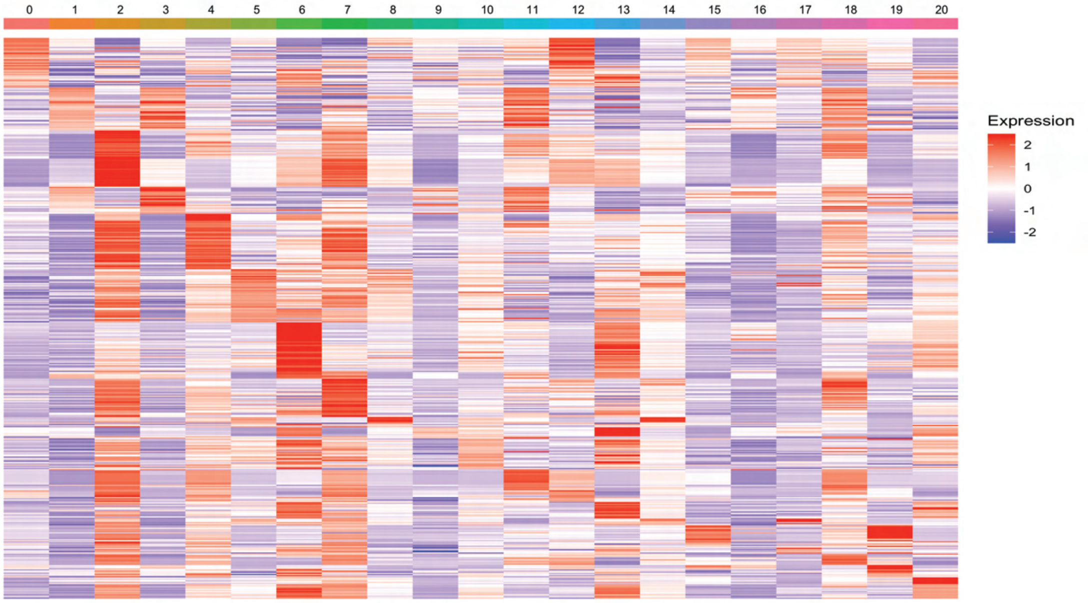


B

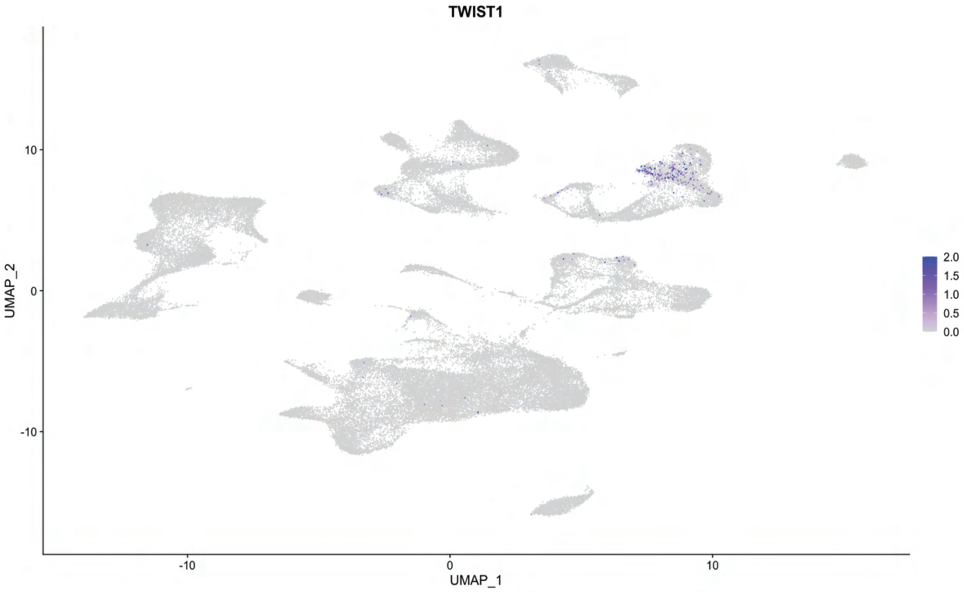


**Supplemental Figure 6.**

Heatmap of top enriched transcription factor motifs by snATAC-seq cluster. Color scale represents a z-score of the average motif activity within each cluster. B) *TWIST1* gene expression in the scRNA-seq dataset depicting fibroblast specific expression.

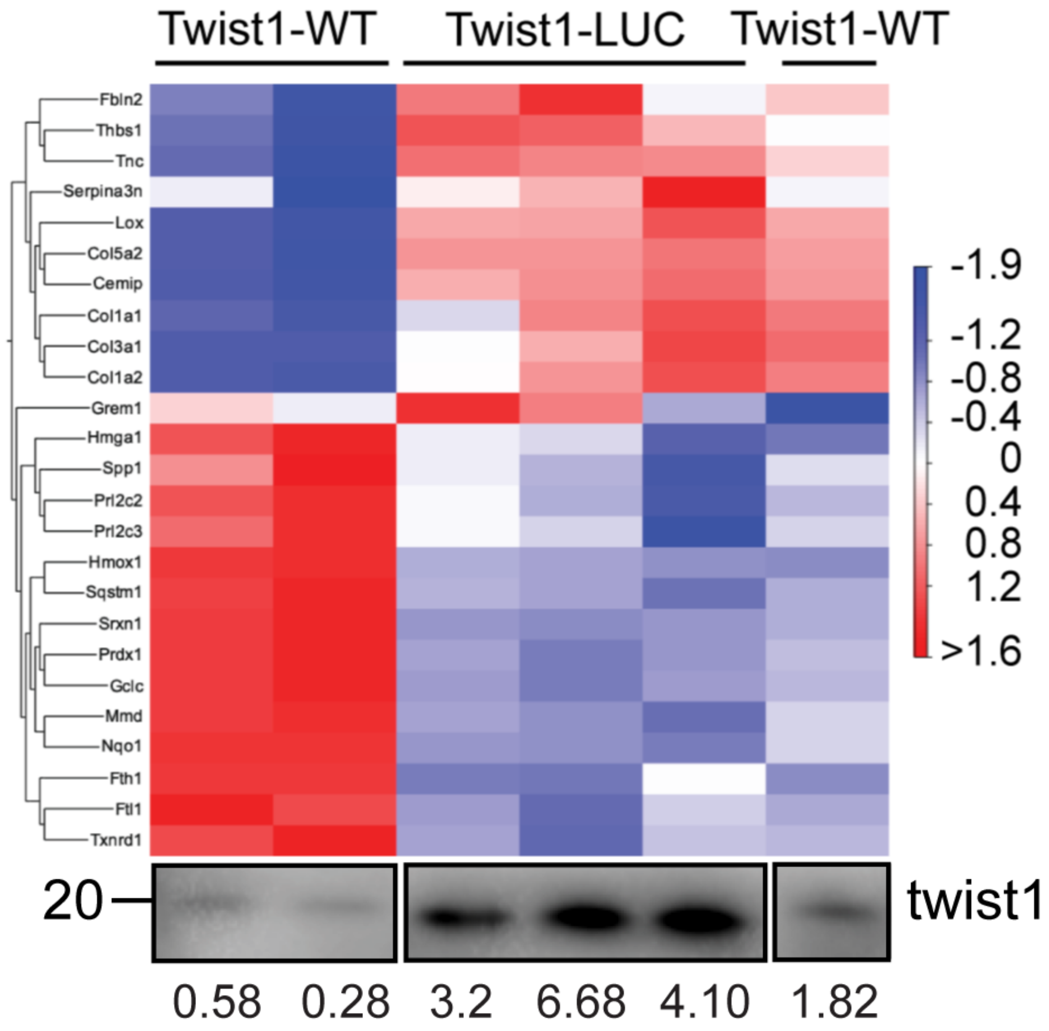


**Supplemental Figure 7**

Gene expression of Twist1-WT and Twist1-LUC murine fibroblasts, with corresponding twist1 immunoprecipitation blot for each sample depicted below. Heatmap was generated using fixed number of features option where given number of features with the highest coefficient of variation (the ratio of the standard deviation to the mean) were kept.

A

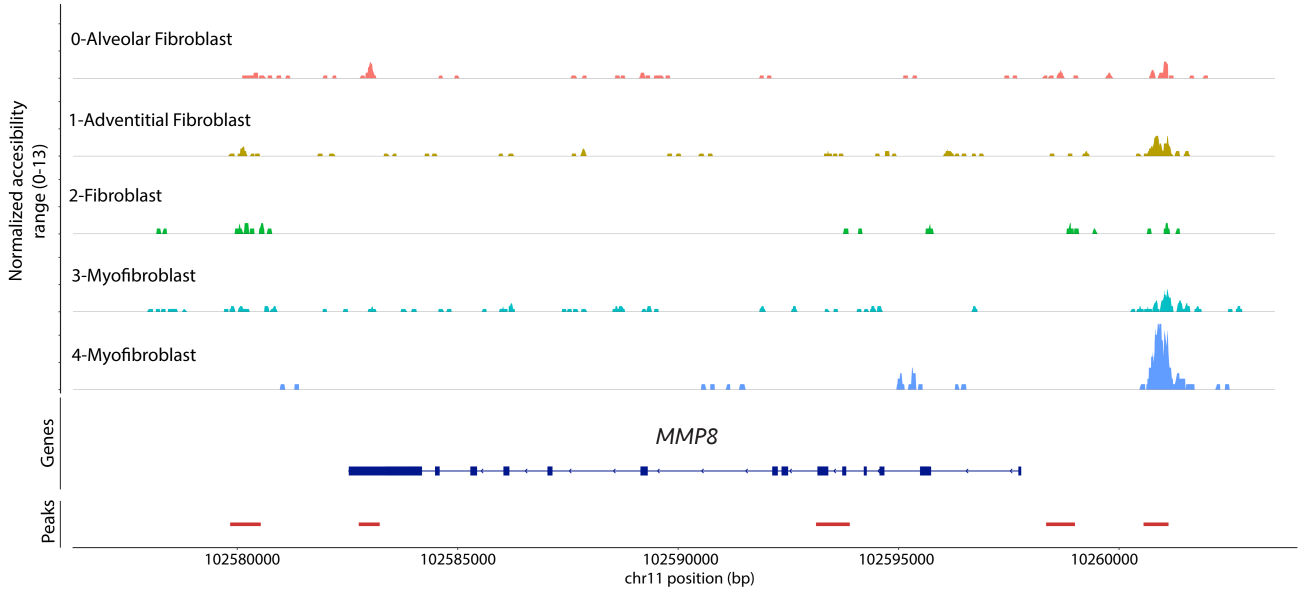


B

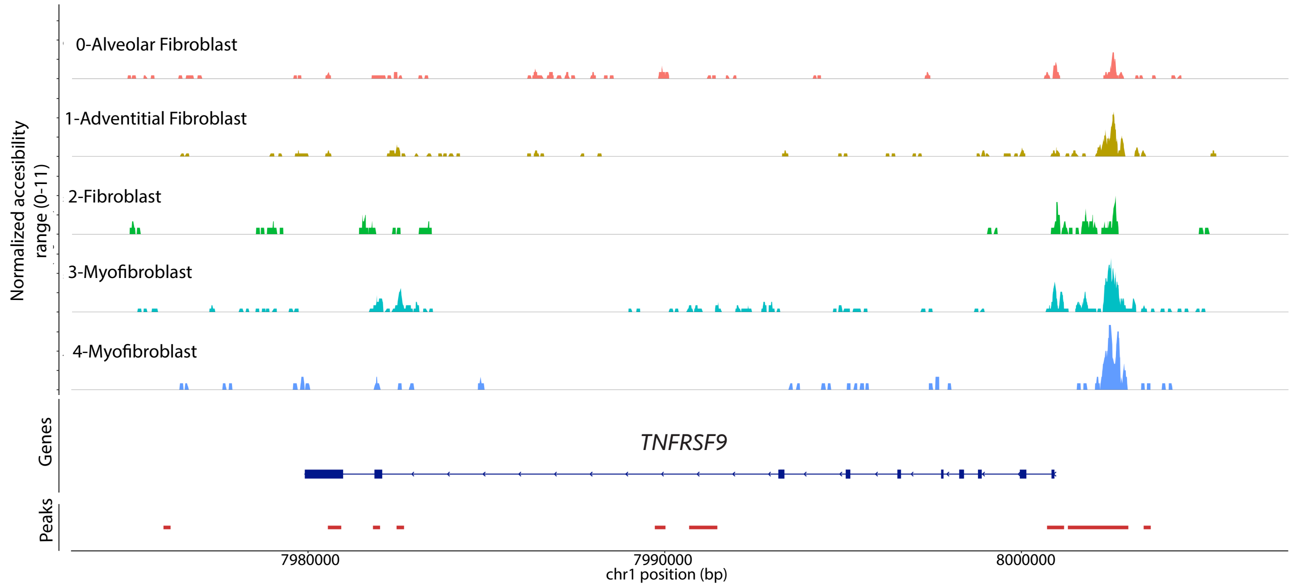


**Supplemental Figure 8**

A) Coverage plot demonstrating Tn5 insertion frequency by snATAC-seq fibroblast cluster in the *MMP8* gene region, with increased chromatin accessibility in the myofibroblasts. B) Coverage plot demonstrating Tn5 insertion frequency by snATAC-seq fibroblast cluster in the *TNFRSF9* gene region, with increased chromatin accessibility in the myofibroblasts.

**Supplemental Table 1**
Demographics for snATAC-seq samples

**Supplemental Table 2**

Demographics for scRNA-seq samples

**Supplemental Table 3**
Top 50 enriched motifs by snATAC-seq cluster

**Supplemental Table 4**
Differentially accessible regions for IPF myofibroblasts vs IPF non-myogenic fibroblasts

**Supplemental Table 5**
Differentially accessible regions for IPF fibroblasts vs control fibroblasts

**Supplemental Table 6**
Differentially expressed genes for IPF myofibroblasts vs IPF non-myogenic fibroblasts

**Supplemental Table 7**
Differentially expressed genes for IPF fibroblasts vs control fibroblasts
